# The Phylogenetic Structure of *β*-diversity: Covariance Matrix Sparsification of Critical Beta-splitting Trees

**DOI:** 10.64898/2026.02.10.705081

**Authors:** Sean P. Svihla, Manuel E. Lladser

## Abstract

Haar-like wavelets sparsify the phylogenetic covariance matrices of large, uniformly random ***k***-regular trees with overwhelmingly high probability. This motivates the Haar-like distance, a ***β***-diversity metric that implicitly ranks the splits of a reference phylogeny by their relevance in differentiating two microbial environments, offering an interpretation as to why the environments differ compositionally. Nevertheless, uniform binary trees exhibit statistical features distinct from those of the trees used by practitioners, leaving the extent of sparsification and the practical validity of the implied Haar-like distance speculative. To address this, our manuscript examines the sparsification of phylogenetic covariance matrices of large critical beta-splitting random trees, a model introduced to better reflect real-world phylogenies. By obtaining sharp asymptotic estimates of the first and second moments of the external path length in this ensemble, we demonstrate that the Haar-like basis also pseudo-diagonalizes the phylogenetic covariance matrix of most large trees in this more realistic framework. Additionally, we devise a test to assess the statistical significance of splits in the reference phylogeny identified by the Haar-like distance. We apply the test to a well-studied microbial mat to further substantiate the presumption that the identified splits represent genuine biological signals differentiating the top and bottom layers of the mat.

## 1 Introduction

In fields such as microbiology, phylogenetic trees arise as models of inter-species covariance (Fukuyama et al. 2012; Pavoine et al. 2004). This prior phylogenetic structure is incorporated into the definition of notable *β*-diversity metrics, which quantify difference between environments in terms of species composition. For example, UniFrac (Lozupone and Knight 2005; Lozupone et al. 2011) uses the notion of shared branch length between samples to capture the relative abundance of observed organisms; on the other hand, DPCoA (Pavoine et al. 2013) directly incorporates the induced covariance matrix of the phylogenetic tree. Phylogenetic *β*-diversity metrics, and especially UniFrac, have had great success in meaningfully ordinating microbial environments based only on their relative abundances of species (Lozupone et al. 2007; Pavoine et al. 2004); however, as newer and more sophisticated reference phylogenies grow larger in size, such metrics face two notable challenges:

1. Phylogenetic covariance matrices are often fully or almost fully dense, so that for large datasets their manipulation is a computational challenge;
2. Traditional phylogenetic *β*-diversity metrics do not provide any indication of *why* two microbial environments are different, limiting their usefulness as exploratory tools.

Fortunately, phylogenetic covariance matrices bear a surprisingly rich structure due to redundancies induced by their hierarchies—in fact, they exactly comprise a set of highly structured matrices called *ultrametric*—and this structure may be exploited to make computations tenable (Gorman and Lladser 2023).

Ultrametric matrices arise in probability theory and statistical physics, among other fields, and, under mild assumptions, arise as covariance matrices of phylogenetic trees. As shown in Nabben and Varga (1994), a matrix *C* is ultrametric if and only if there exists a rooted binary tree *T* = (*V, E*) and function *ℓ* : *E* → ℝ_+_ such that

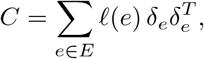

where *δ*_*e*_ is a binary column vector with entries indexed by the leaves of *T*, indicating those that descend from an edge *e*. This identity is equivalent to

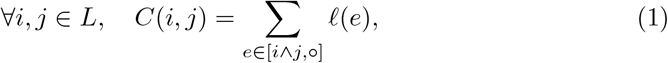

where *L* is the leaf set of *T* and [*i* ∧ *j*, ◦] denotes the set of edges connecting (*i* ∧ *j*), the least common ancestor of leaves *i* and *j*, with the root of the tree, denoted ◦. In the setting of rooted trees, a matrix with entries such as in (1) is called a *phylogenetic covariance matrix*.

The formulation of phylogenetic covariance in (1) is equivalent to a model of neutral evolution wherein the average value of a trait in a population follows a Brownian motion (e.g., (Hansen and Martins 1996; Harmon 2019)). This is a reasonable model for phylogenetic trees of microbial communities, which are commonly assembled by clustering 16s rRNA (Woese 1987; Tringe and Hugenholtz 2008). Under this model, trait variation between lineages evolves as independent Brownian motions from the time the lineages diverge, and accordingly the covariance in (1) is their shared path length to the root—a proxy for time before evolutionary divergence. Notably, this formulation matches one’s intuition that species which shared more of their evolutionary history should thrive or struggle similarly in similar environments.

Recent work has demonstrated that the phylogenetic covariance matrix associated with a large, uniformly-at-random binary (Gorman and Lladser 2023) or more generally *k*-regular (Svihla and Lladser 2024) tree is pseudo-diagonalized (i.e., significantly sparsified) with high probability after changing basis to the so-called Haar-like wavelets (Gavish et al. 2010) of the tree. In particular, the fraction of non-zero off-diagonal entries in the transformed matrix converges in probability to zero as the size of the tree approaches infinity. Such sparsification enables computational manipulation of the covariance matrix, which would otherwise be infeasible. For instance, in Gorman and Lladser (2023) the Haar-like basis was used to sparsify the covariance matrix of the 97% Greengenes tree, a widespread reference phylogeny in earlier metagenomic studies (DeSantis et al. 2006; McDonald et al. 2012). The sparsification also motivated the introduction of a new *β*-diversity metric, the Haar-like distance (Gorman and Lladser 2023), which unlike its predecessors may highlight the mutations—and consequently the biological functions—that best explain the differences in the microbial relative abundance between two environments. More recently, using the same reference phylogeny, Gorman and Lladser (2024) showed that a kernel derived from Haar-like wavelets can be optimized against a random forest in order to learn a *β*-diversity metric, the adapted Haar-like distance, that preserves much of the random forest’s accuracy but renders its discrimination rules interpretable in terms of splits along the tree.

This degree of sparsification results from the multiscale nature of the Haar-like wavelets and the phylogenetic covariance matrix structure, and it requires that the tree be shaped appropriately (in a sense that will be specified below) to be effective. Not all families of trees are so well behaved (for example, the phylogenetic covariance matrices of binary caterpillar trees are not sparsified by the Haar-like wavelets). The results in Gorman and Lladser (2023); Svihla and Lladser (2024) depend on the characteristics of the uniform binary tree ensemble. As a consequence, inasmuch as the uniform ensemble falls short of modeling real-world phylogenies, such results may not generalize beyond the 97% Greengenes tree. In fact, Aldous (2001) observed that the median size of the smaller subtree after a split in trees with *n* leaves scales as O (*n*^1*/*2^) in real-world phylogenies, whereas under a uniform distribution it scales as O (log *n*). Because tree balance is related to subtree split size and plays a key role in sparsification under the Haar-like basis, this fact brings into question whether the wavelets will be as effective at sparsifying the covariance matrices of large trees used by practitioners.

Our manuscript considers a more realistic random tree model, the critical beta-splitting random tree, introduced by Aldous (1996) as one better suited for real-world phylogenies (see also Aldous (2001)). After a detailed asymptotic analysis of the first and second moments of the external path length in a large random tree from this ensemble—complementing recent results in Aldous and Janson (2024a,b) and extending some of Aldous and Pittel (2023); Aldous and Janson (2024c)—we show that the Haar-like basis pseudo-diagonalizes with overwhelmingly high probability the phylogenetic covariance matrix of large critical beta-splitting random trees. In particular, the effects of the Haar-like wavelets on the phylogenetic convariance matrix of the 97% Greengenes tree may not have been merely coincidental and could generalize to a much wider range of phylogenies, including new and more detailed ones being developed for metagenomic studies such as Greengenes2 (McDonald et al. 2024), SILVA (Quast et al. 2013; Yilmaz et al. 2014), and WoL (Zhu et al. 2019).

Further, we build upon the analysis in Gorman and Lladser (2023) of the Guerrero Negro microbial mat samples in Kirk Harris et al. (2013) by devising a hypothesis test with which to assess the statistical significance of dominant components of the Haar-like distance. Importantly, the results further substantiate the claim that dominant components represent genuine biological signal that is not explainable by sampling variance or other random effects.

In the remainder of the introduction, we briefly introduce some notation and terminology and provide an overview of concepts and properties relevant for presenting and proving our new results.

### 1.1 Notation and terminology

Throughout the manuscript, we regard functions from *L* to ℝ interchangeably as vectors. 1_*A*_(*x*) denotes the indicator function over set *A*.

We use standard terminology for trees. We consider planted, ordered, and unlabelled trees. In particular, *T* = (*V, E*) is a rooted tree with vertex set *V* and edge (branch) set *E*. We denote its root by ◦. Because *T* is planted, ◦ has degree one, i.e., only one edge is connected to it. The sets of leaves and interior nodes of *T* are denoted by *L* and *I*, respectively, and the size of the tree is |*T* | := |*L*|, the number of leaves. By convention, we label the leaves of the tree *i* = 1, 2, …, |*T* | in the order they are encountered in its postorder traversal (i.e., their left-to-right order). See Figure 1 for an illustration.

**Fig. 1:**
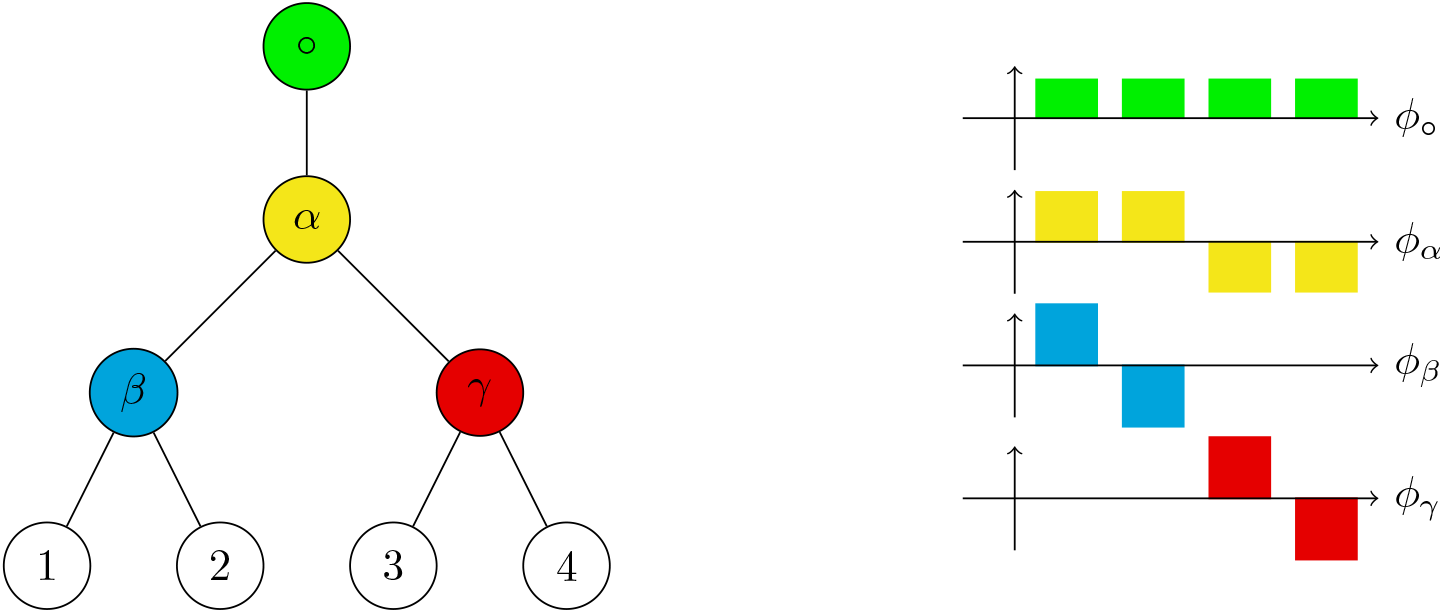
An illustration of a planted binary tree and associated Haar-like wavelets. Left: A balanced binary tree with leaves labelled 1, 2, 3, 4 and interior nodes labelled *α, β, γ*. Note that the trees we consider need not be balanced. Right: The Haar-like wavelets associated with the tree on the left. Each wavelet is associated with a split in the tree and defined to be positive over the leaves of the left subtree and negative over those of the right.

The depth of a *v* ∈ *V*, denoted depth (*v*), is its edge-distance to the root. The *external path length* of *T* is the quantity

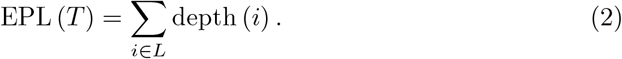

Each tree is equipped with an edge length function *ℓ* : *E* → ℝ_+_. Following Gorman and Lladser (2023), the *trace length* of *T* is the function *ℓ*^*^ : *E* → ℝ_+_ defined as

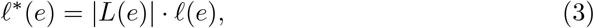

where *L*(*e*) are the set of leaves that descend from an edge *e*. (Likewise, define *L*(*v*) as the set of leaves that descend from a node *v*.) For *u, v* ∈ *V*, define

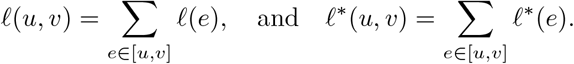

We denote by *C* the phylogenetic covariance matrix of *T* in the standard basis. We denote the Haar-like basis matrix by Φ. (The precise definitions of these are given in the next section.)

### 1.2 The Haar-like wavelets

We first define Haar-like wavelets associated with a planted binary tree. See Gavish et al. (2010) for details regarding the construction of the wavelets, and Gorman and Lladser (2023); Svihla and Lladser (2024) for their use in the context of phylogenetic trees.

#### Definition 1.1.

*(Gorman and Lladser 2023) The Haar-like basis associated with the tree T is the set* {*φ*_*v*_}_*v*∈*I*_ *defined as follows:*

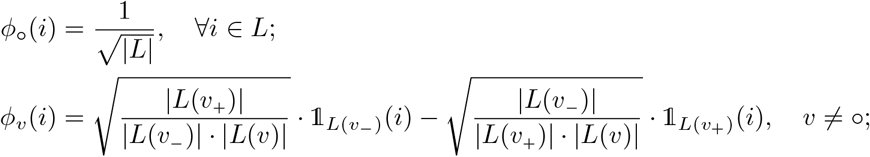

*where v*_−_ *and v*_+_ *denote the left and right children of v, respectively*.

*The functions φ*_*v*_, *with v* ∈ *I, are mutually orthogonal with respect to the standard inner-product in* ℝ^|*L*|^, *and form a basis for the linear space of functions from L to* ℝ. *Let* Φ *denote the Haar-like basis matrix (i*.*e*., *the matrix with columns φ*_*v*_ *for v* ∈ *I)*.

It was shown in Gorman and Lladser (2023, Theorem 2.3) for binary trees and as a corollary to Svihla and Lladser (2024, Theorem 2) for *k*-regular trees that

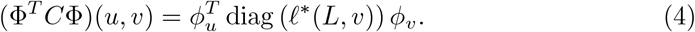

As a consequence, (Φ^*T*^ *C*Φ)(*u, v*) = 0 whenever *φ*_*u*_ and *φ*_*v*_ have disjoint supports. By counting such interactions, one obtains the following (Gorman and Lladser 2023, Corollary 3.5):

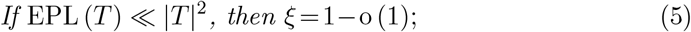

where *ξ* denotes the proportion of non-zero, off-diagonal entries, in Φ^*T*^ *C*Φ. In particular, Gorman and Lladser (2023, Theorem 3.4) gives the bound

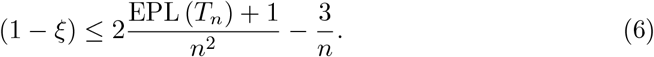

If the condition in (5) is satisfied, Φ^*T*^ *C*Φ is pseudo-diagonal, i.e., nearly all of its off-diagonal entries are zero. This property is the key reason why wavelets may significantly reduce computations when working with tree phylogenetic covariance matrices.

### 1.3 The critical beta-splitting random tree

The critical beta-splitting ensemble is a special case of a one-parameter family of random binary trees introduced by Aldous (1996), called beta-splitting models. These models generate a tree by a random, recursive binary partitioning of [*n*] according to a splitting probability distribution *q*(*n, i*), *i* = 1, …, *n* − 1, which determines the probability that [*n*] is split into clades of size *i* and (*n* − *i*), respectively. The critical beta-splitting tree model uses

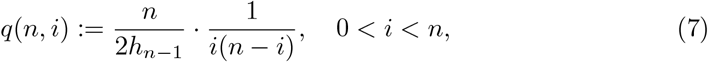

where *h*_*n*−1_ denotes the (*n* − 1) th harmonic number, 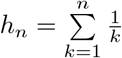 for *n* ≥ 1.

Our goal in further analyzing the critical beta-splitting tree is to obtain similar sparsification results to Gorman and Lladser (2023); Svihla and Lladser (2024) but for a more realistic model of real-world phylogenetic trees than the uniform model used therein. To do so, we require accurate estimates of the first and second moments of the external path length in a large, random critical beta-splitting tree. Our analysis builds on recent results and techniques about critical beta-splitting tree model from recent works by Aldous and Pittel (2023); Aldous and Janson (2024a,b,c).

As noted in Aldous (2001), the critical beta-splitting tree is a more suitable model in at least one important respect: *in many real-world phylogenetic datasets, among clades of n leaves, the median size of the smaller sub-clade scales as* 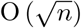. (See top left of Figure 2.) The critical beta-splitting tree shares this property, whereas traditional random models typically have a median minor sub-clade size that scales as O (log *n*) or Θ (*n*) (Aldous 1996, 2001). Because the arguments in Gorman and Lladser (2023); Svihla and Lladser (2024) rely on the statistics of uniformly random trees, from which the hypothesis in (5) follows, it is unclear whether sparsification extends to the critical beta-splitting ensemble. In fact, as shown in the bottom panel of Figure 2, uniform binary trees exhibit median minor-subtree-split sizes that differ markedly from those of the widely used Greengenes trees. This contrasts with the close similarity in median values observed between Greengenes trees and critical beta-splitting trees of various sizes, as shown in the top-right panel of Figure 2. The rate of decay of this ratio appears to be faster for critical beta-splitting trees than for the uniform ensemble, implying that this ensemble exhibits a higher degree of sparsity for the same sized tree than is the uniform ensemble.

**Fig. 2:**
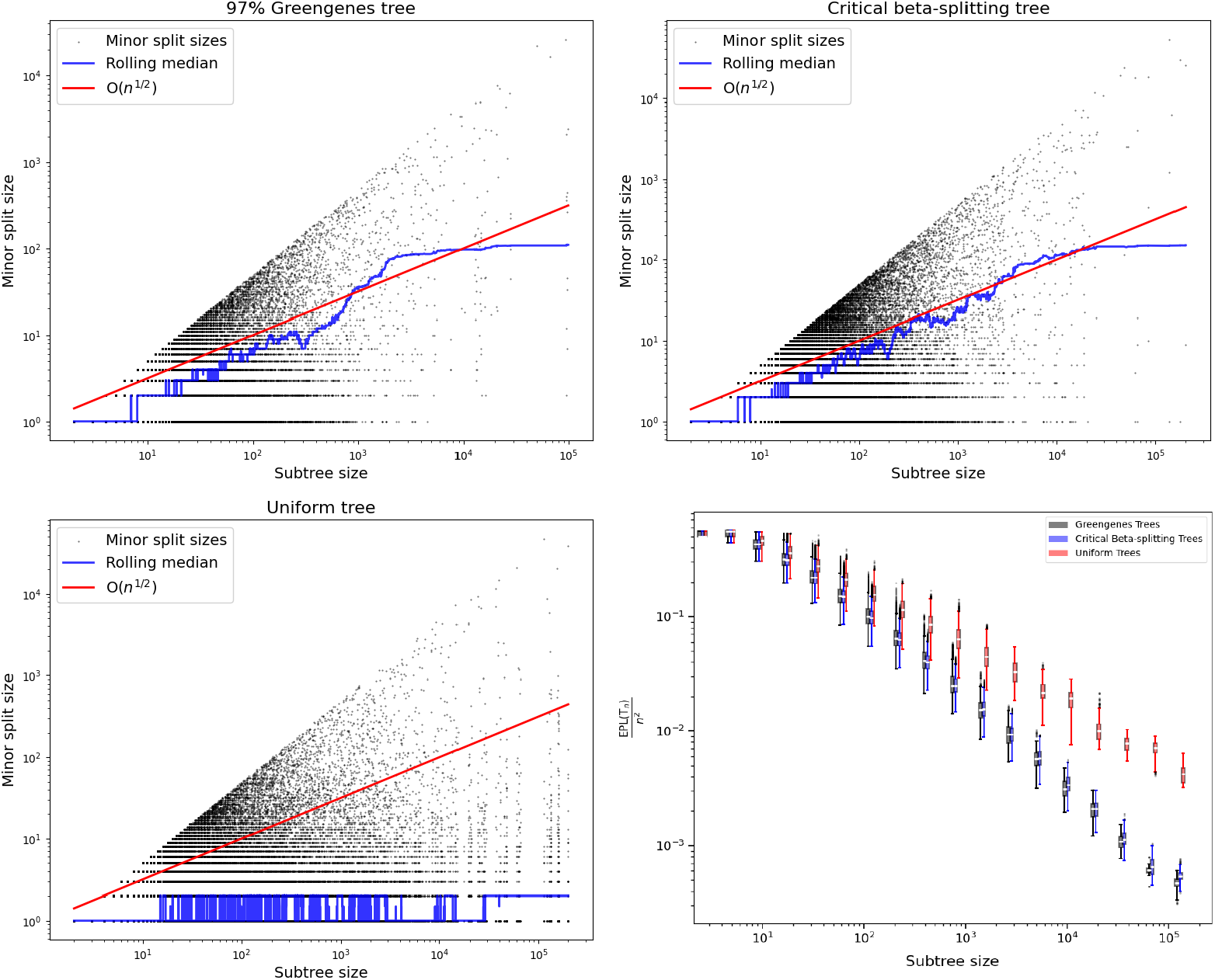
Comparison of median minor subtree size and the ratio EPL (T_n_) */*n^2^ of various families of binary trees. Subfigures (a)-(c): Median minor split size is shown for the 97% Greengenes tree, critical beta-splitting random tree, and uniform random tree, respectively. Figures (b) and (c) depict a single realization of the respective random tree. We observe that the median minor split size of the critical beta splitting tree grows as O (*n*^1*/*2^) like that of the 97$ Greengenes tree, while that of the uniform tree does not. Subfigure (c): The ratio EPL (T_**n**_) */***n**^**2**^ is shown for varying tree size *n* for the 97% Greengenes tree, critical beta-splitting tree, and uniform tree. Box plots in the figure were computed from *N* = 30 samples for each displayed subtree size. We again observe that the critical beta-splitting tree better aligns with the qualitative behavior of the real-world 97% Greengenes tree.

**Fig. 3:**
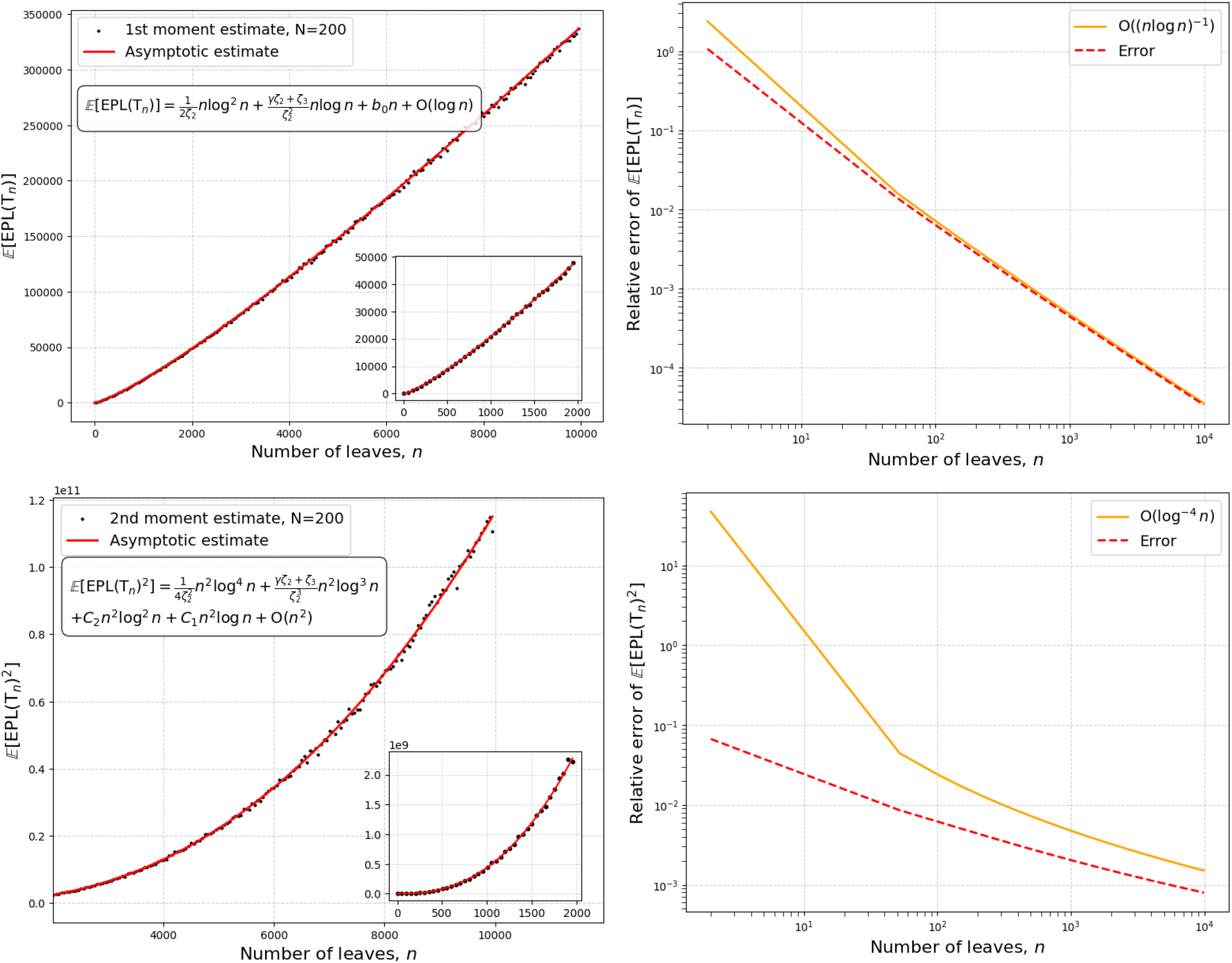
Comparison of asymptotic estimates of moments with numerically computed exact values. Plots of the first moment estimate (top left), first moment relative error (top right), second moment estimate (bottom left), and second moment relative error (bottom right). Exact values were computed numerically from the recurrence relations in Aldous and Pittel (2023) and (23). We see that the asymptotic estimates of the first two moments coincide with sample moments estimated from *N* = 200 draws of the critical beta-splitting distribution. The relative errors of these estimates are consistent with the constants of our derived expansions being accurate, the errors for the first and second moment estimates being bounded by O (*n* log(*n*))^−1^ and O (log^−4^(*n*)), respectively.

In what follows, T_*n*_ denotes a critical beta-splitting random tree with *n* leaves. To extend the results in Gorman and Lladser (2023); Svihla and Lladser (2024) to this new setting, we are interested in the random variable *L*_*n*_, defined to be the edge-depth (i.e., the number of edges from a node to the root) of a leaf chosen uniformly at random from T_*n*_.

## 2 Haar-like Wavelets Sparsify Realistic Phylogenies

The Haar-like wavelets are capable of sparsifying the covariance matrix of a large phylogenetic tree—irrespective of the distribution from which the tree was drawn— provided that its topology is suitable to cancellation of terms. This result was established in Gorman and Lladser (2023, Corollary 3.5) and, more generally, in Svihla and Lladser (2024, Corollary 5), using the result in (5). What remains to be seen is how often EPL (*T*_*n*_) ≪ *n*^2^ when *n* is large.

To this end, let *L*_*n,i*_, with *i* = 1, …, *n*, denote the edge-depths of the leaves in T_*n*_. The random vector 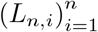 is exchangeable; in particular, ∀*i* ∈ [*n*], *L*_*n,i*_ ∼ *L*_*n*_, the edge-depth of a leaf selected uniformly at random. The external path length of T_*n*_ may, then, be written in terms of these edge-depths as

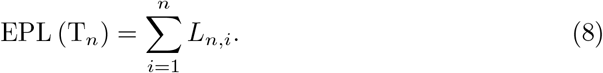

The next result characterizes asymptotically the first and second moments of EPL (T_*n*_). Its proof and some auxiliary results can be found in Section 5.

### Theorem 2.1.

*Let* T_*n*_ *be a random tree with n leaves, distributed according to the critical beta-splitting model. Then, the following asymptotic estimates hold:*

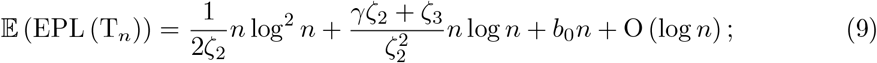

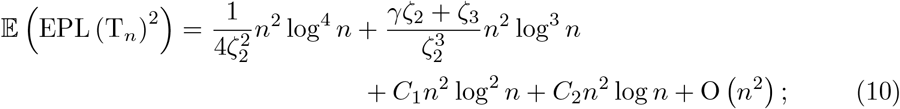

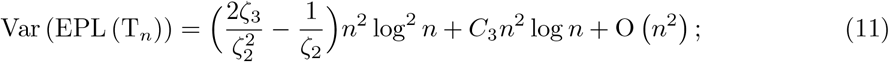

*where*

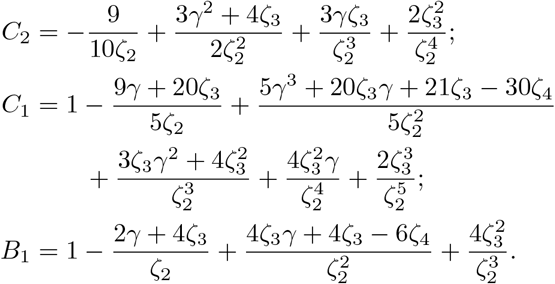

We note that the asymptotic estimate for 𝔼 (EPL (T_*n*_)) in Theorem 2.1 is similar to the expansion for 𝔼 (*L*_*n*_) in (14) due to 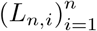 being exchangeable. In addition, 𝔼 (EPL (T_*n*_))^2^ and 𝔼 (EPL (T_*n*_))^2^ have identical leading terms, so that they cancel out when estimating Var (EPL (T_*n*_)).

Our next result extends Gorman and Lladser (2023, Corollary 3.8) from the uniform binary tree model to the critical beta-splitting tree model. It demonstrates that the Haar-like wavelets are remarkably effective at sparsifying the phylogenetic covariance matrix of large binary trees generated under the more realistic critical beta-splitting model. Its proof is omitted as it mirrors those of Gorman and Lladser (2023, Corollary 3.8) and Svihla and Lladser (2024, Corollary 15). We provide a fast implementation of the sparsification algorithm on GitHub (see our Data Availability Statement in Section 4).

### Corollary 2.2.

*Let* T_*n*_ *be a tree with n leaves, distributed according to the critical beta-splitting model, and C*_*n*_ *be its covariance matrix. If* Φ_*n*_ *is the Haar-like matrix associated with* T_*n*_, *and ξ*_*n*_ *the fraction of vanishing entries in* 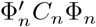, *then* 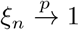 *as n* → ∞.

## 3 Statistical Significance of Haar-components

The Haar-like wavelets motivate the definition of the Haar-like distance (Gorman and Lladser 2023), a phylogenetic *β*-diversity metric between microbial environments. This metric is notable because it readily points to splits in the reference tree that contribute most and least to the distance between any pair of environments, a feature lacking in traditional *β*-diversity metrics.

Samples of microbial environments are represented by their counts of operational taxonomic units (OTUs). These are obtained by mapping DNA reads from a sequenced environment to the leaves of *T*. Accordingly, we represent microbial samples as functions from *L* to the nonnegative integers. So, if *a* : *L* →ℕ then *a*_*i*_, with *i* ∈ *L*, denotes the total number of reads designated as OTU *i* in sample *a*. We define |*a*| := ∑_*i*∈*L*_ *a*_*i*_ as the total OTU count in the sample; in particular, *a/*|*a*| represents the relative abundance of OTUs in sample *a*.

### Definition 3.1

(Haar-like Distance and Components (Gorman and Lladser 2023)). *The Haar-like distance between two microbial samples a, b* : *L* →ℕ *is the quantity*

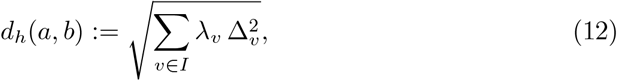

*where* 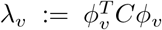 *is the v-th entry in the diagonal of the matrix* Φ^*T*^ *C*Φ; *and* 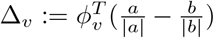 *is the v-th coordinate of the vector* 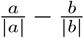*in the Haar-like basis*.

*We call λ*_*v*_ 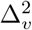, *with v* ∈ *I, a Haar-component of the distance between a and b*.

Because each Haar-component in (12) corresponds to a unique interior node in *T*, the magnitude of the Haar-like distance between two samples *a* and *b* can be explained in terms of the individual splits in the tree: *If λ*_*v*_ 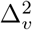 *is large (small) relative to the other components, then v may play a (not) significant role in differentiating the environments from which the samples were collected*. Accordingly, the largest Haar-components may elucidate the mutations—and hence biological adaptations—that are most relevant for distinguishing *a* from *b* by their OTU compositions. Nevertheless, no test has been proposed in the literature to assess the statistical significance of comparatively large Haar-components. In the remainder of this section, we show how to assess this significance using a permutation test over the OTU counts descending from a split within microbial samples *a* and *b*. Our approach makes no strong assumptions about the underlying distribution of OTUs in the environments. As a proof-of-concept, we revisit the statistical significance of splits in the 97% Greengenes tree identified in Gorman and Lladser (2023) as key differentiators between the shallowest and deepest samples from a microbial mat (Kirk Harris et al. 2013).

### 3.1 Methodology

Let *a* and *b* be samples from two microbial environments. For an interior node *v* in *T* and a threshold *t >* 0, we aim to assess the statistical significance of *λ*_*v*_ 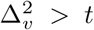 following a permutation test principle (e.g., (Evans and Matsen 2012)).

Under the null hypothesis that the samples come from compositionally identical environments, the DNA reads are exchangeable across them. In particular, we may estimate the cumulative distribution function (c.d.f.) of *λ*_*v*_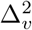 by resampling *a*, drawing without replacement |*a*| of the |*a*|+|*b*| OTUs, and reassigning the unselected OTUs to *b*. After each resample, *λ*_*v*_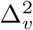 is recomputed from the resampled vectors of OTU counts from each environment; these recomputed values are then used to estimate empirically the c.d.f. of *λ*_*v*_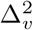. The p-value reported to assess the statistical significance of the Haar-component associated with *v* in *d*_*h*_(*a, b*) is the empirical probability that *λ*_*v*_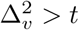.

We note that according to the Dvoretzky-Kiefer-Wolfowitz (DKW) inequality (Dvoretzky et al. 1956; Massart 1990), estimating the c.d.f. within an infinity-norm error of *ϵ*, with probability (1 − *δ*), requires *N* = log(2*/δ*)*/*(2 *ϵ* ^2^) resamples.

In practice, we resample *a* by drawing the OTU counts from a multivariate hypergeometric distribution with parameters (|*a*|; *a*+*b*) (Johnson 2005; Johnson et al. 1997), rather than by brute-force permutations. We provide fast hypergeometric and multivariate hypergeometric samplers on GitHub (see our Data Availability Statement in Section 4), which implements the algorithm described in Hörmann (1994) based on the rejection-inversion sampler for the tail of a log-concave distribution described in Hörmann and Derflinger (1996). This package is used to define an estimator for the c.d.f. of 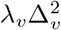 under the null hypothesis. Further, it is implemented so that no significant additional computational expense is incurred by estimating the c.d.f.s of multiple Haar-components at once, nor by estimating one or more c.d.f.s at multiple points.

While the dominant Haar-components help identify the *most* relevant splits in terms of the Haar-like distance, we are not limited to considering only these components. By filtering Haar-components with the permutation test, one is able to distinguish genuine environmental differences from random effects at finer scales, where it would be impossible to deduce from the Haar-component magnitudes alone. This can be accomplished by defining

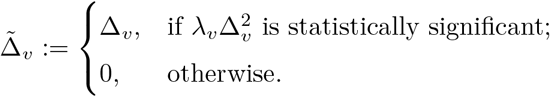

Then 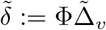, is the projection of 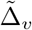 back to the standard basis, a proxy for (*a* − *b*). Each of these Haar-components represents a dimension along which the observed environmental differences cannot be explained by chance; consequently, they constitute a (relatively) low-dimensional subspace in which to represent and study these environments. In practice, since there are many non-zero Haar-components, we assess the statistical significance after applying a Bonferroni correction. See Figure 6 for an example of this in practice.

**Fig. 4:**
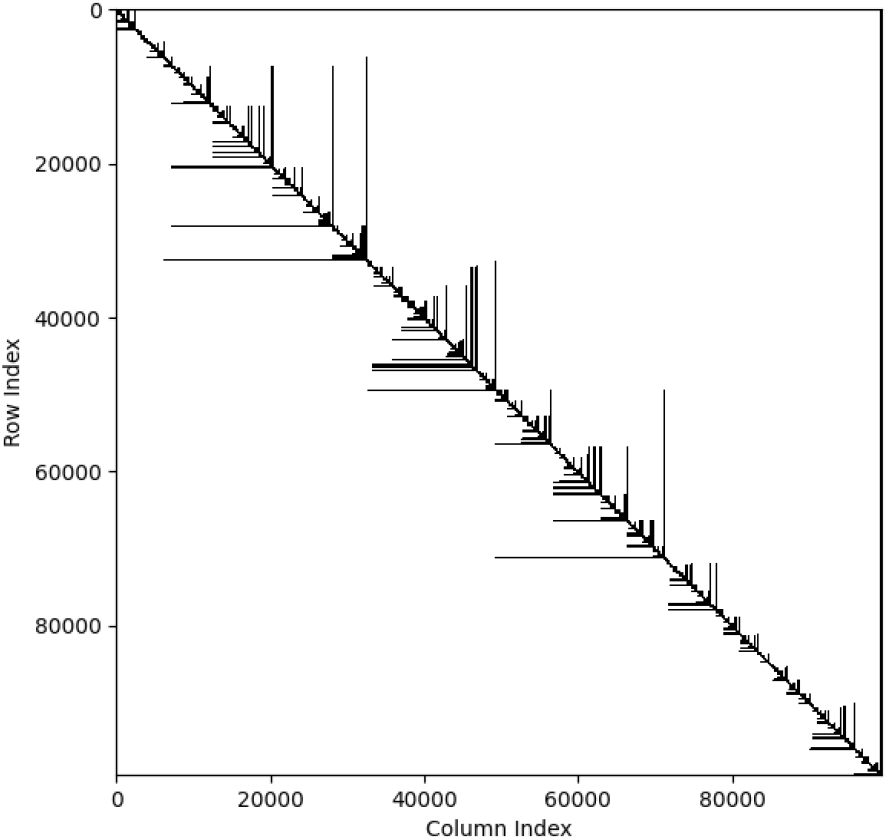
Sparsity pattern of the covariance matrix of 97% Greengenes tree in the Haar-like basis. The 97% Greengenes tree has |*L*| = 99, 322 leaves, for a total of 9, 864, 859, 684 entries in its dense phylogenetic covariance matrix. We see that the resulting sparse matrix is pseudo-diagonalized, with an overwhelming proportion of off-diagonal entries being zero. If stored as an array of double precision floats, the dense matrix would be ≈74 GB; the sparse matrix, by contrast, is ≈76 MB. The bound in (6) guarantees a sparsify of at least 99.8992% off-diagonal entries vanishing. The true sparsity is 99.8993%.

**Fig. 5:**
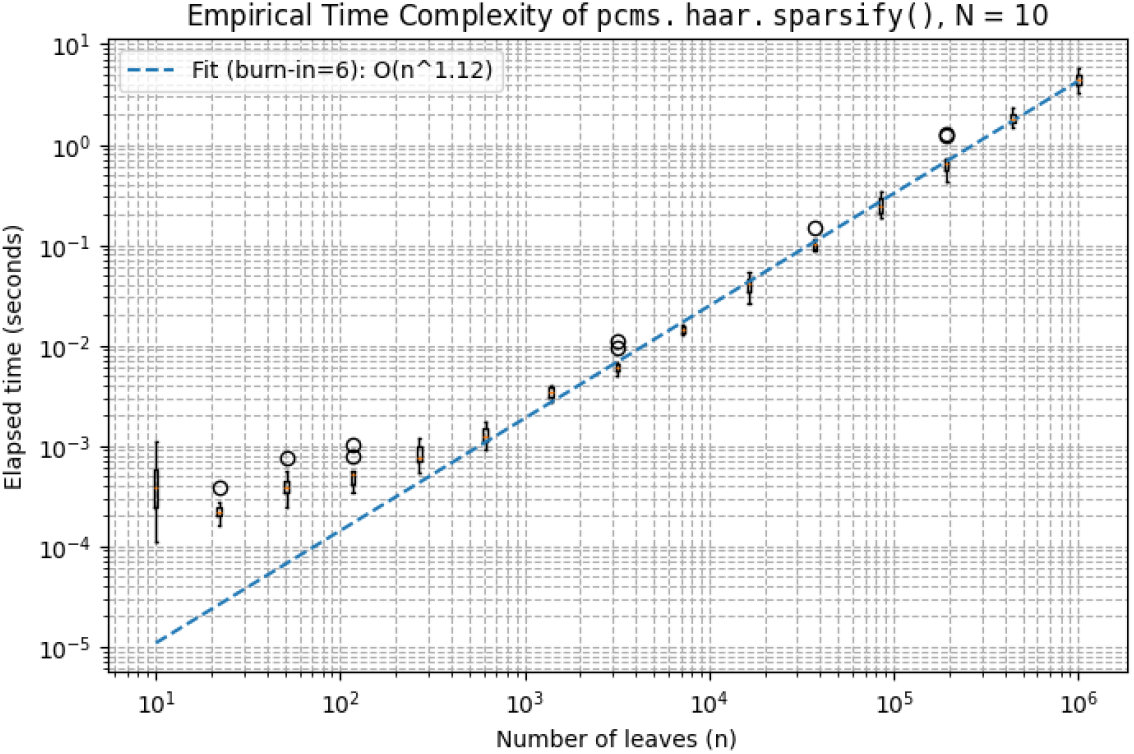
Empirical Time Complexity of Sparsification Algorithm. The empirical time-complexity is sub-quadratic in the number of leaves of large trees.

**Fig. 6:**
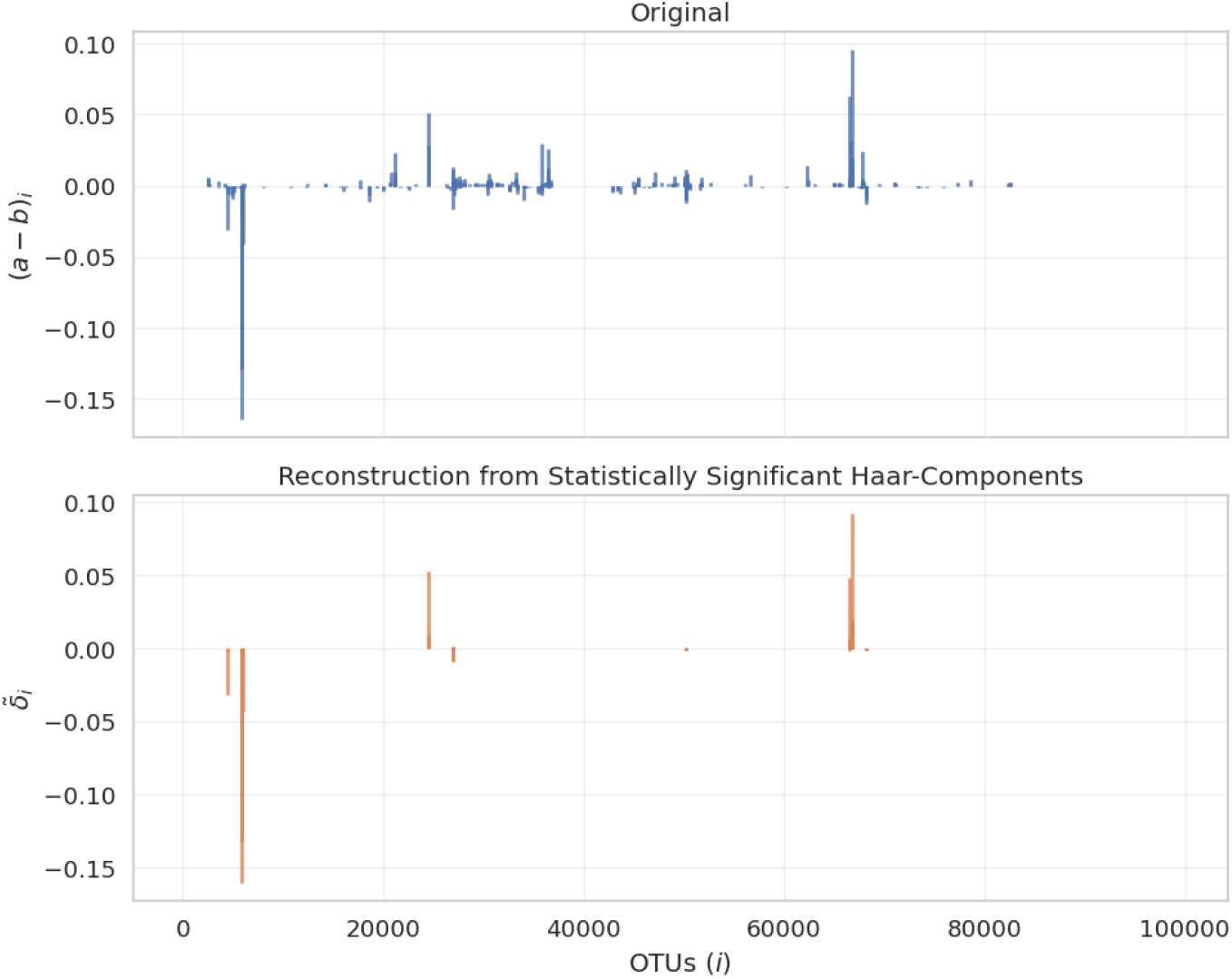
Reconstruction of relative OTU counts of the 2013 Guerrero Negro study from only statistically significant Haar-components. Significance was assessed using a Bonferroni correction on the raw *p*-values for *m* = 4271 hypotheses, the number of non-zero Haar components. As shown in the bottom plot, 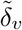, a proxy for (*a*−*b*), retains its dominant features with only 159 statistically significant coordinates.

### 3.2 Proof-of-concept: the Guerrero Negro microbial mat

Microbial mats are multi-layered biofilms of microbial communities, typically only a few centimeters thick at most, often fostering a rich biodiversity and exhibiting sharp variation in species composition across space and time (Dillon et al. 2009; Hörnlein et al. 2018). The Guerrero Negro microbial mat, located in Baja California Sur, Mexico, is one of the most well-studied mats (Ley et al. 2006; Lozupone and Knight 2007; Dillon et al. 2009; Maza-Márquez et al. 2024). In Kirk Harris et al. (2013), multiple samples from a single mat at Guerrero Negro were collected, sectioned into horizontal layers (at different depths reflecting the vertical stratification of the mat), and pooled within each layer. In this section, we assess the statistical significance of the twelve largest Haar-components differentiating the shallowest and deepest samples in this 2013 study.

Since the samples in this study were collected concurrently, from the same pond, and nearby one another within a narrow radius, it is reasonable to assume that the OTU composition at each level is homogeneous. In particular, the pooled reads from each layer may be regarded as i.i.d. draws from a single probability distribution. This allows us to use the test in Section 3.1 to assess the statistical significance of the components of the Haar-like distance between samples from the top (0–1 mm) and bottom (34–49 mm) layers. We computed the Haar-like distance between the top and bottom samples using the OTU counts published to Qiita (study ID 1200), derived from closed-reference clustering of 454 reads against the 97% Greengenes tree (Kirk Harris et al. 2013; McDonald et al. 2012). Out of 4271 non-zero Haarcomponents, 956 had a raw p-value less than 0.05 and 159 remained statistically significant after applying a Bonferroni correction.

In the case of this study, the Haar-like distance yields a wealth of subtle (yet nonetheless significant) relationships to explore between the top- and bottom-level environments. However, under the assumption that the largest Haar-components represent the most important splits (inasmuch as they contribute most to the magnitude of *d*_*h*_(*a, b*)), we primarily concern ourselves with the several most dominant components. We will find support for this assumption when we observe that many of the compositional differences identified in Kirk Harris et al. (2013) are manifest in a small subset of dominant Haar-components.

The twelve most dominant Haar-components are displayed in Figure 7, with nodes indexed in postorder traversal. Table 1 summarizes the findings of the permutation test. For all but one of the twelve dominant Haar-like components considered, the magnitude of *λ*_*v*_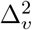 is statistically significant at the 5% level under the null hypothesis that the top- and bottom-level environments of the Guerrero Negro microbial mat are compositionally indistinguishable. The three most dominant Haar-components have also been deemed significant using a different criterion (Gorman and Lladser 2023).

**Table 1:**
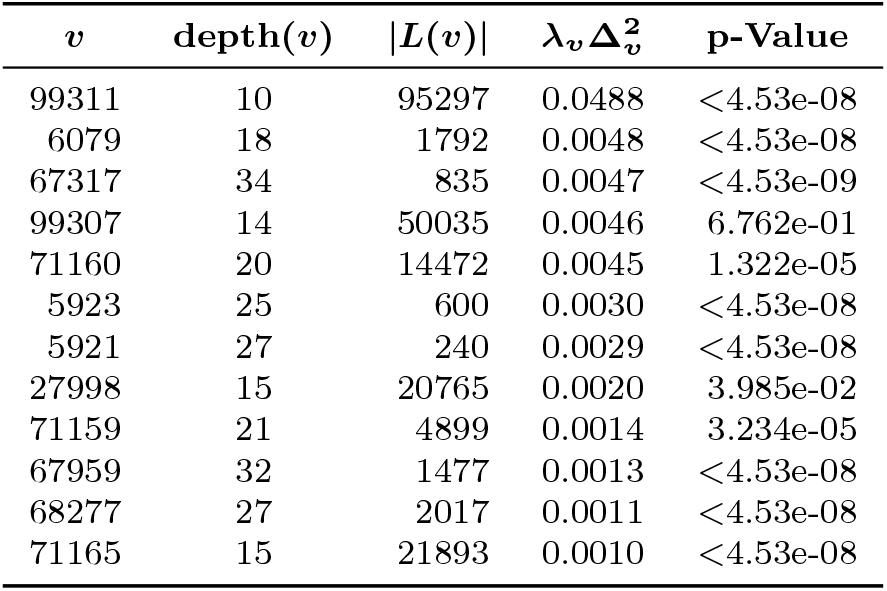
Statistical significance of the twelve most dominant Haar-components differentiating the shallowest and deepest samples in the 2013 Guerrero Negro study. We obtained p-values from the null empirical c.d.f. of *λ*_*v*_ 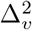 for each interior node *v* in the 97% Greengenes tree, using *N* resamples chosen according to the DKW inequality with *E* = 0.0001 and *δ* = 0.01 (see Section 3.1). Where the permutation algorithm failed to sample sufficiently far into the tail, returning zero, we report the discovery probability based on *N*. The reported p-values include a Bonferroni correction for *m* = 12 hypotheses. For reference, the 97% Greengenes tree has a median leaf depth of 51 and a maximum leaf depth of 107. Components corresponding to deeper nodes capture finer-scale information. *Remark: The Haar-component associated with v* = 99311 *was misreported in Gorman and Lladser (2023) but is corrected here*.

**Fig. 7:**
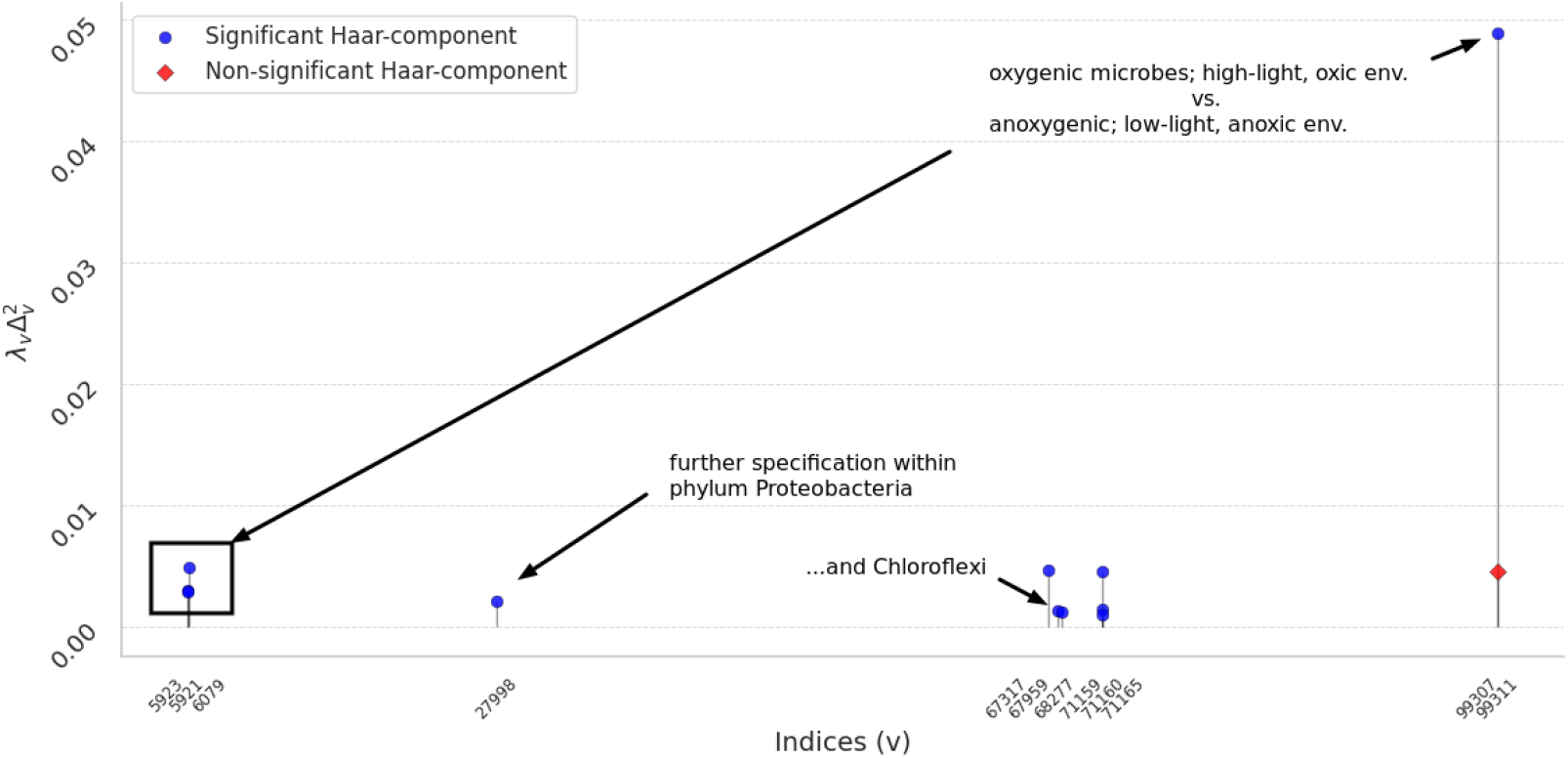
Plot of the twelve largest components of the Haar-like distance between top- and bottom-level 2013 samples from the Guerrero Negro microbial mat. Each point is marked according to whether the null hypothesis that these top and bottom samples are compositionally indistinguishable is rejected at the 5% significance level. We see that all but one of the top twelve dominant Haar-components pictured are statistically significant under our test. Each of these Haar-like components is readily interpretable in terms of splits in the reference phylogeny; our annotations include some potentially interesting interpretations of the dominant splits.

Figure 8 illustrates some of the dominant Haar-components in terms of the taxa found beneath their corresponding interior nodes. The most dominant component at node 99311 marks a split characterized by a prevalence of Cyanobacteria in the top layer relative to other bacterial phyla (such as Chloroflexi and Proteobacteria) in the bottom layer, as previously noted in Gorman and Lladser (2023). Nodes 6079, 5923, and 5921 mark splits exhibiting different responses to increasing mat depth within the phylum Cyanobacteria and, further, within the class Oscillatoriophycideae. Similarly, nodes 71159, 71160, and 71165 appear to capture the prevalence of the phylum Chloroflexi over the other bacterial phyla marked off from Cyanobacteria by node 99311. In turn, node 68277 marks a different response by the class Anaerolineae as compared with other members of Chloroflexi, with nodes 67317 and 67959 further delineating distinct responses within Anaerolineae. Finally, node 27998 marks a split within Proteobacteria revealing an increase in Alphaproteobacteria and a decrease in Betaproteobacteria and Gammaproteobacteria between the top and bottom environments.

**Fig. 8:**
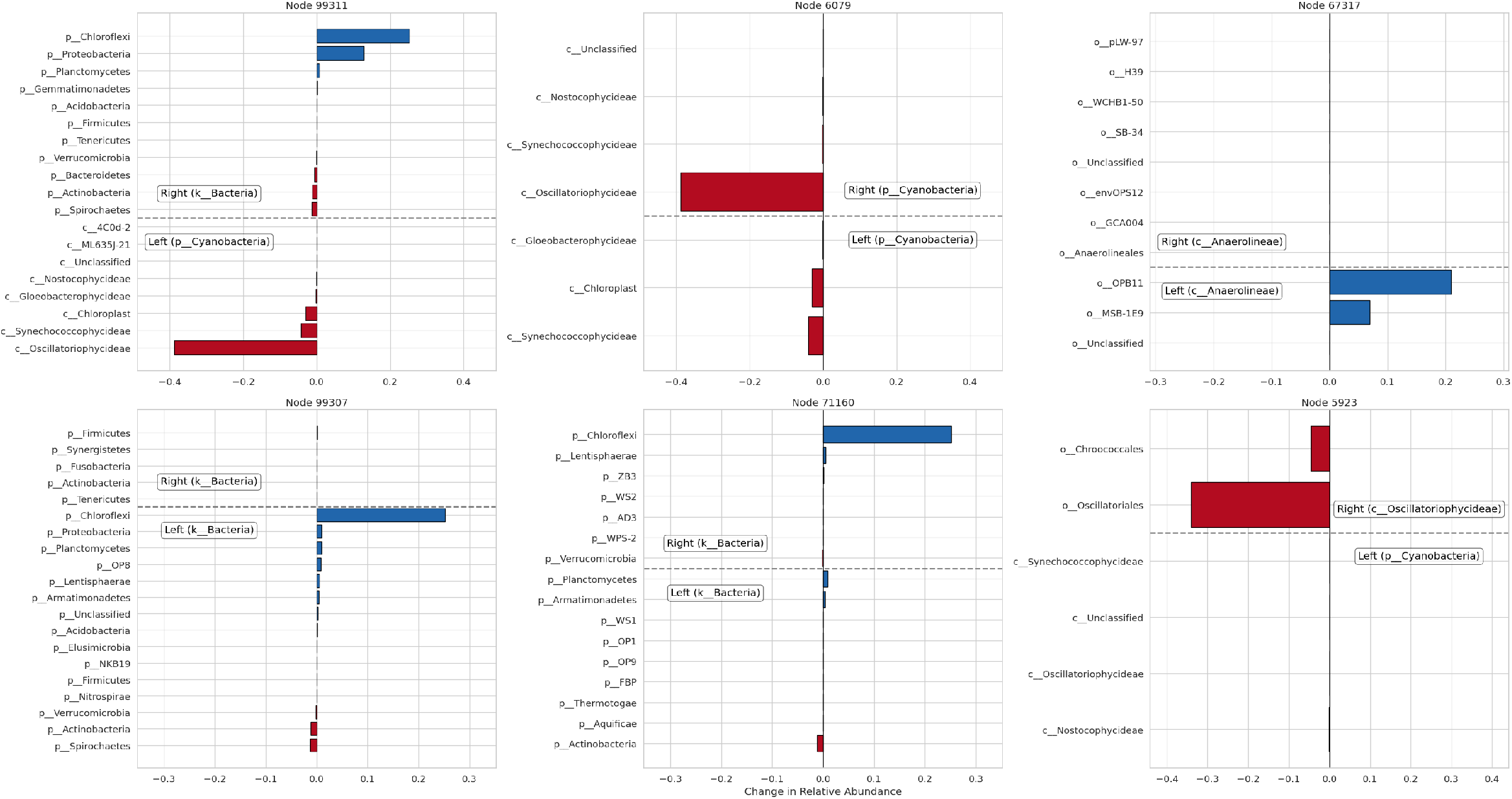
Relative taxonomic composition breakdown between top and shallow samples from Guerrero Negro. Differences in relative abundance of OTUs between the deep layer (*a*) and the shallow layer (*b*) of the Guerrero Negro microbial mat beneath the splits of the six most dominant Haar-components. A positive *x*-value indicates prevalence in *a* over *b*, and a negative value indicates prevalence in *b* over *a*. Plots are ordered by descending value of the Haar-components from left to right and top to bottom. For each split *v*, the text box gives the lowest-level taxonomic rank shared by all OTUs in the respective subtree. Further, each OTU, *i*, beneath *v* is grouped into a class *I*_*k*_ by its taxonomic classification at the next lowest rank. The value reported is the net change in relative abundance within each of these classes (i.e., 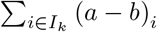).

The dominant Haar-components suggest a gradient between the top and bottom layers marked by a shift from an oxic, high-light environment populated by oxygenic phototrophs to an anoxic, low-light environment populated by anoxygenic phototrophs (Kurth et al. 2021; Kawai et al. 2021). Such a spatial gradient has been widely observed and studied in microbial mats (Ley et al. 2006; Lozupone and Knight 2007; Dillon et al. 2009; Kirk Harris et al. 2013; McDonald et al. 2012; Hörnlein et al. 2018; Maza-Márquez et al. 2024). It is noteworthy, then, that the Haar-like distance implicitly identifies a low-dimensional space, the (Haar-like) coordinates of which are directly interpretable in terms of such a gradient.

Unlike the other twelve most dominant Haar-components, node 99307, associated with the fourth dominant component, is not statistically significant at the 5% level of the permutation test. It was noted in Gorman and Lladser (2023) that *λ*_*v*_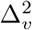 depends on the difference between the averages of *f* = (*a/*|*a*| − *b/*|*b*|) on the leaves of the left- and right-subtrees of an interior node *v*. Upon inspection, however, one finds that the top and bottom layers are composed overwhelmingly of OTUs from the left-subtree, although their compositions within that subtree differ. The comparatively large magnitude of *λ*_99307_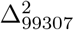 is therefore not due to a difference in OTU composition between environments across the split, but rather to a prevalence of OTUs from one subtree in both environments, contrasted with a shared lack of abundance in the other. In other words, the split does not correspond to a compositional difference between environments; rather, both are characterized by a prevalence of left-subtree OTUs. The permutation test is robust to such artifacts, and the Haar-component appears non-significant accordingly.

## 4 Final Remarks

Phylogenetically informed *β*-diversity metrics such as UniFrac and DPCoA have found widespread use and success in metagenomics. DPCoA is defined in terms of a covariance matrix derived from a reference phylogeny and often leads to qualitatively similar conclusions to UniFrac. This manuscript is motivated by two shortcomings of such metrics: (i) the covariance matrix is often large and dense, making computational manipulation is expensive and potentially infeasible; and (ii) these metrics are not readily interpretable in terms of the phylogeny.

Previous work has shown that the Haar-like basis and distance can address these concerns, ultimately due to the tendency of the Haar-like basis to significantly sparsify the phylogenetic covariance matrix of large phylogenetic trees chosen uniformly at random. However, because the significant sparsification relies heavily on the statistical properties of these random tree models, it was unclear whether it holds for large phylogenetic trees with features consistent with those used by practitioners. We have addressed this issue by deriving sparsification results similar to those but under the more realistic critical beta-splitting model of random binary trees. In particular, by obtaining sharp asymptotic estimates of the first and second moments of the external path length of the trees in this ensemble, we have shown that the Haar-like basis is highly effective at sparsifying all but a negligible fraction of large trees produced by this model. It remains an open problem to determine the rate at which EPL (T_*n*_) */n*^2^ converges to 0 under the critical beta-splitting model.

In addition to enabling the sparsification of large, dense phylogenetic covariance matrices, the Haar-like basis motivates the definition of the Haar-like distance, a *β*-diversity metric which importantly provides an implicit ranking of the splits in the reference phylogeny by order of their relevance in differentiating between two microbial environments. Still, it is not obvious that these dominant coordinates successfully parse genuine signal from noise. To address this uncertainty, we devised a permutation test based on the null hypothesis that a pair of environmental samples were drawn from two environments which are compositionally indistinguishable. The result of this test allows one to conclude when the magnitude of a particular Haar-like distance component is not plausibly explainable by random effects between samples from identical environments.

In the proof-of-concept considered here, we found that all but one of the twelve most dominant Haar-like components were highly statistically significant, and that these components described many of the macro-level compositional-environmental differences identified in the original study. In particular, the dominant Haar-components suggested a gradient between the top and bottom layers of the Guerrero Negro microbial mat which is characterized by a shift from an oxic, high-light environment populated by oxygenic phototrophs to an anoxic, low-light environment populated by anoxygenic phototrophs—a spatial gradient which is well-known and widely observed in microbial mats like those at Guerrero Negro. While we have chosen a well-studied environment as proof-of-concept, t, this behavior supports the use of the Haar-like distance as an exploratory tool in understudied environments, where dominant Haar components may reveal unknown gradients.

Our analysis shows that the technique of sparsifying dense phylogenetic covariance matrices by a change-of-basis to the Haar-like wavelets is well-justified under the critical beta-splitting model. The fact that we obtain similar sparsification-in-probability results under this more realistic model suggests that the previously observed sparsification on the 97% Greengenes tree was not coincidental, but rather reflects a general behavior of real-world phylogenetic covariance matrices with respect to the Haar-like basis. Furthermore, our demonstration of a permutation test on samples from the Guerrero Negro microbial mat substantiates the claim that dominant components capture genuine biological signals, while offering a method to study subtle—yet statistically significant—differences between environments.

### Data Availability Statement

We have provided a package which implements the phylogenetic covariance matrix sparsification described here at https://github.com/spsvihla/pcms, which also contains Python notebooks with which to reprorduce all figures in this manuscript. We have made use of the 97% Greengenes tree (McDonald et al. 2012) and data from the Kirk Harris et al. (2013) Guerrero Negro microbial mat study. We have provided information on how to access these data in the README file of the GitHub repository.

## 5 Proof of Theorem 2.1

In this section, we present the proof of Theorem 2.1, which is comprised mostly of Lemmas 5.4 and 5.5. Our arguments are overall inspired by those of Aldous and Pittel (2023, Theorem 2.4). In particular, recall that *L*_*n*_ denotes the edge-depth of a leaf selected uniformly at random in *T*_*n*_ and that first moment of *L*_*n*_ obeys the recursion (Aldous and Pittel 2023, (2.41))

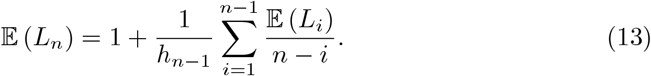

We make use of asymptotic formulas for the mean and variance of *L*_*n*_ (Aldous and Pittel 2023, Theorem 1.2):

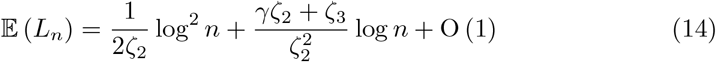

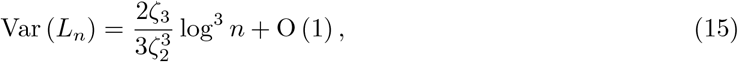

where *ζ*_*s*_, for *s >* 1, is the Riemann-zeta function and *γ* is Euler’s constant. The following refinement of (14) will be particularly useful in our analysis (Aldous and Janson 2024c, Theorem 1.2):

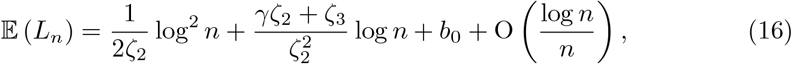

where

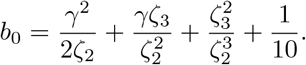

Lastly, we borrow from Aldous and Pittel (2023, Proposition 2.3) the identities

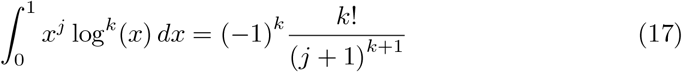

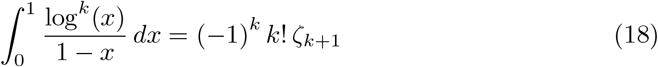

for integers *j* ≥ 0 and *k* ≥ 1.

In addition, we make use the following technical lemmas, the proofs of which can be found in Appendix A.

### Lemma 5.1.

*For all integers k* ≥ 1:

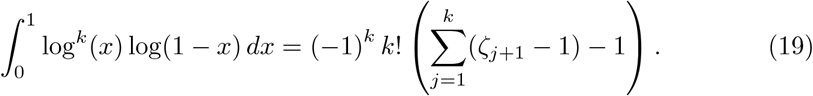

### Lemma 5.2.

*For all integers k* ≥ 1:

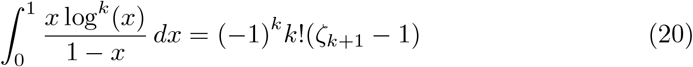

### Lemma 5.3.

*Let c* ∈ ℝ. *Then, uniformly for all integers n > c and m* ≥ 1:

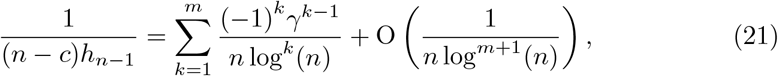

Returning to the proof of Theorem 2.1, recall that *L*_*n,i*_ denotes the edge-depth of the leaf with label *i*, 1 ≤ *i* ≤ *n*. To complete the proof of the theorem, we require two additional results which we state and prove next. Lemma 5.4 derives a recursion for 𝔼 (*L*_*i*_ · *L*_*j*_), and Lemma 5.5 finds an approximate solution to the recursion that is accurate up to O (1).

### Lemma 5.4.

*For i* ≠ *j, the following recursion applies:*

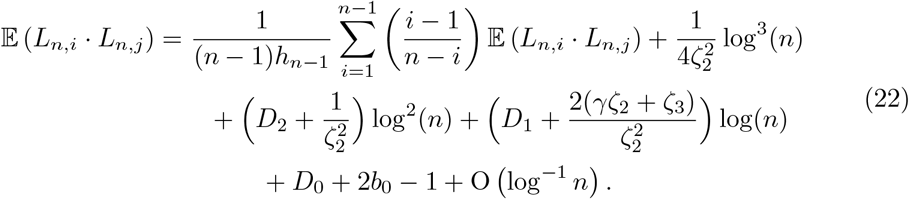

*where*

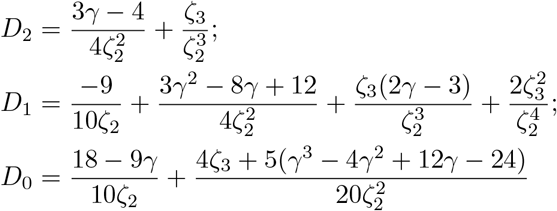

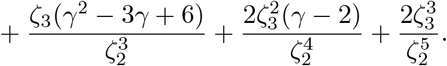

*Proof* Without loss of generality, assume that *i* = 1 and *j* = 2, and define *Z*_*n*_ := *L*_*n*,1_ *· L*_*n*,2_ for brevity. Let *I*_*n*_ be the (random) size of the left subtree of *T*_*n*_ after the first split, so that the right-subtree has size (*n − I*_*n*_). Then, the conditional distribution of *Z*_*n*_ given that *I*_*n*_ = *I* satisfies the distributional recursion

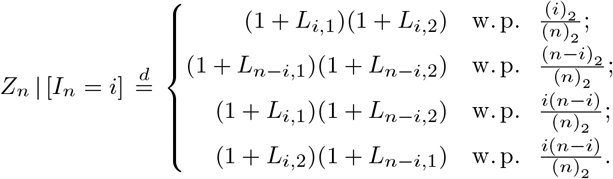

The first two cases correspond to the events in which both leaves fall on the left or right subtree, respectively; the final two cases correspond to these leaves falling into different subtrees, in which case their edge-depths are conditionally independent. So, the conditional expectation of *Z*_*n*_ given the split size is

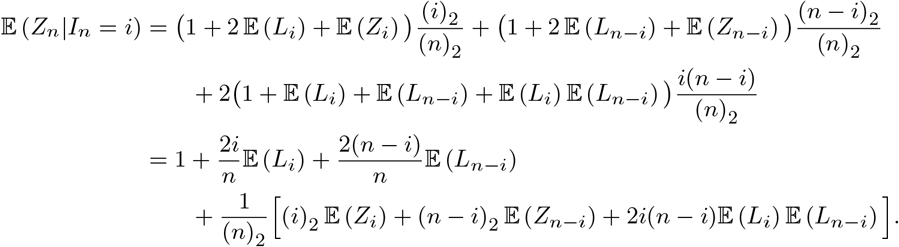

The Law of Total Expectation implies that the expectation of *Z*_*n*_, after some simplification, satisfies

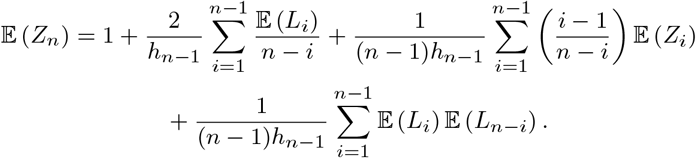

Due to (13), the first two terms on the right hand-side above simplify to 2𝔼 (*L*_*n*_) *−* 1. Thus, we may rewrite the last expression for 𝔼 (*Z*_*n*_) as

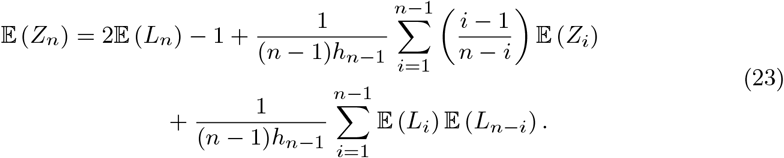

Our task is now to derive asymptotic estimates for the first and last terms in the above recursion. In the remainder of the proof, we demonstrate how to handle the last term only, as the other can be handled similarly.

To begin, substituting (16) into 𝔼 (*L*_*i*_) 𝔼 (*L*_*n*−*i*_), we obtain that

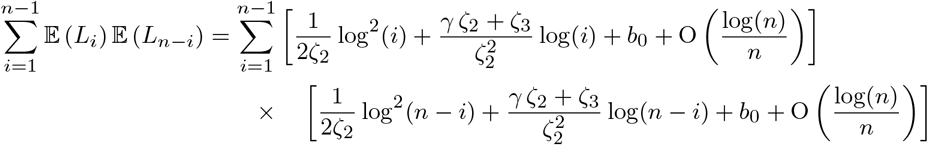

which becomes

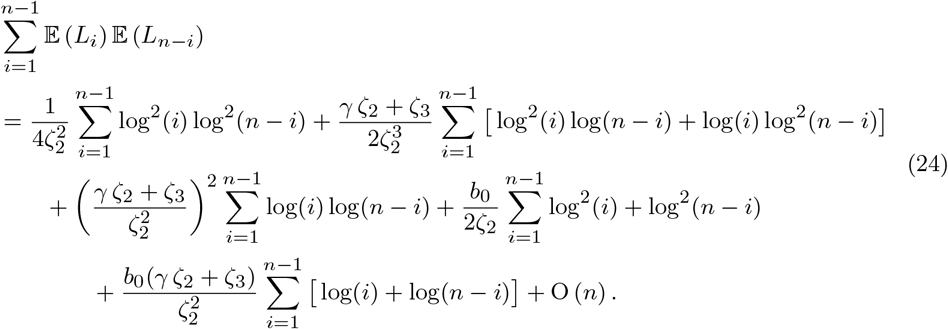

Next, one can obtain asymptotic formulas for each of the summations above by repeatedly applying the following argument, which we only demonstrate for the left-most sum in (24) using Riemann sum arguments. First note that

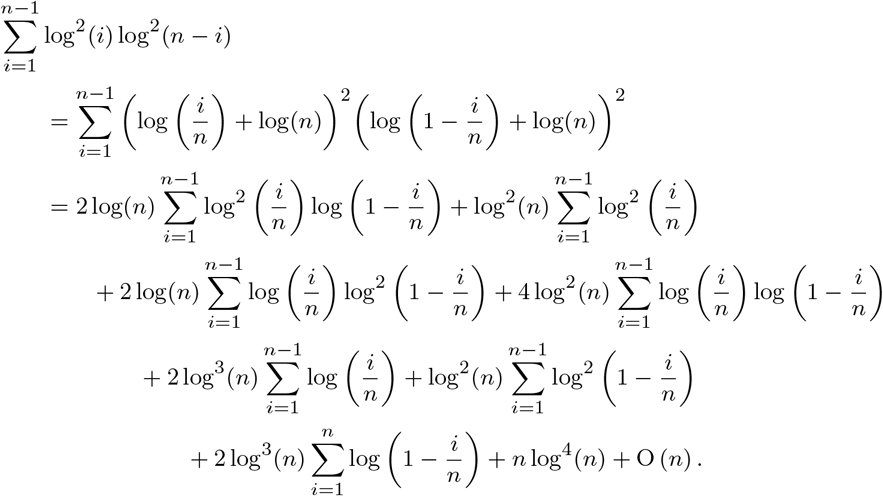

From the identity in (17) and Lemma 5.1, it follows that

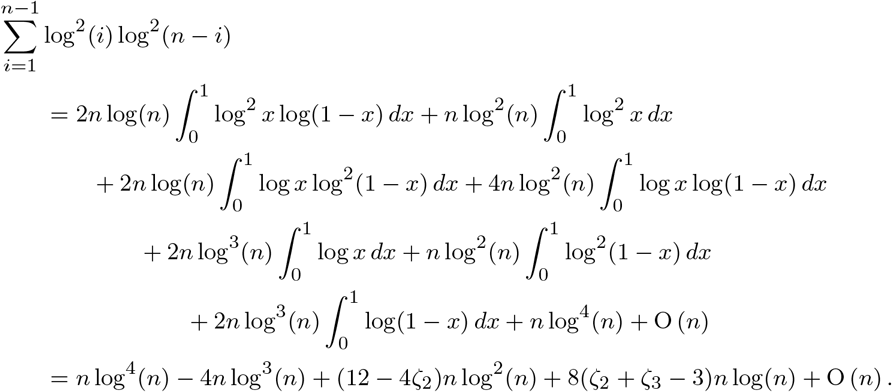

Using similar arguments for the other terms and Lemma 5.3, we obtain that

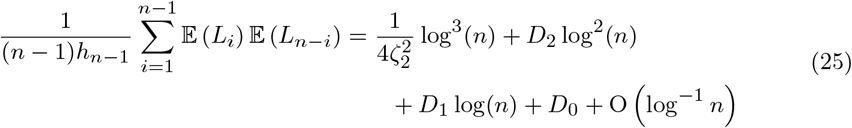

for *D*_0_, *D*_1_, and *D*_2_ as defined in the statement of the lemma. Finally, substituting (25) and (16) into (23), we obtain the implicit recursion for 𝔼 (*Z*_*n*_) in the lemma.

### Lemma 5.5.

*For i* ≠ *j, the following recursion applies:*

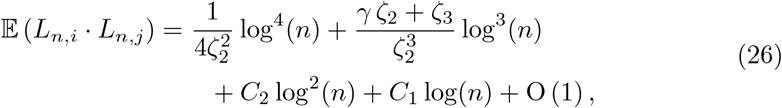

*where*

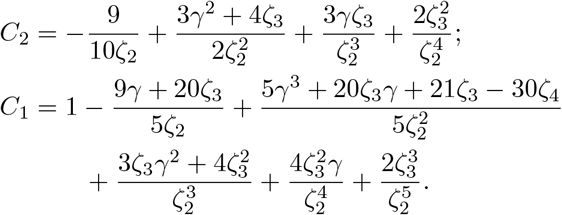

*Proof* We start by noting the following useful identities:

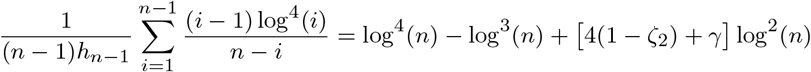

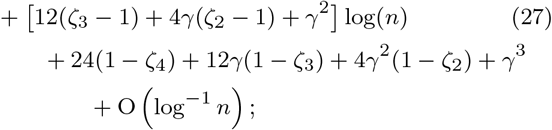

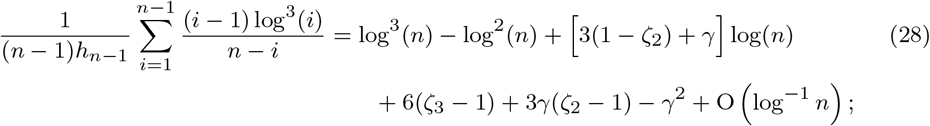

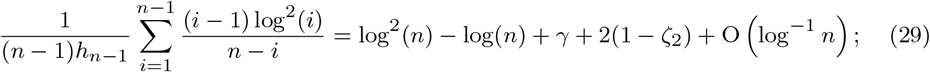

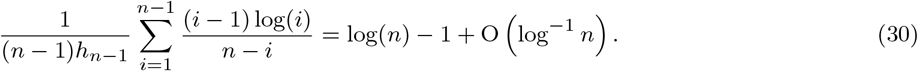

The first two identities follow from 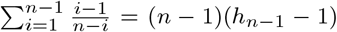 and the application of Lemmas 5.2–5.3. The last two identities can be found in the proof of Aldous and Pittel (2023, Theorem 2.4).

The key idea of the proof is to seek an approximate solution to the recursion in (22) of the form *E*(*n*) = *A* log^4^(*n*) + *B* log^3^(*n*) + *C* log^2^(*n*) + *D* log(*n*) for suitable constants *A, B, C*, and *D*, and then show that 𝔼 ( *L*_*n,i*_ *· L*_*n,j*_) = *E*(*n*) + O (1), as in Aldous and Pittel (2023).

By substituting *E*(*n*) for 𝔼 (*Z*_*i*_) in (22), we find that a sum remains to be handled. In particular, the first term becomes

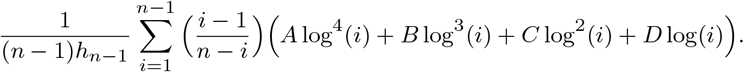

Considering each power of log(*i*) in turn, we find that (27)-(30) provide an asymptotic estimate for this sum, which constrains the constants *A, B, C*, and *D* to satisfy:

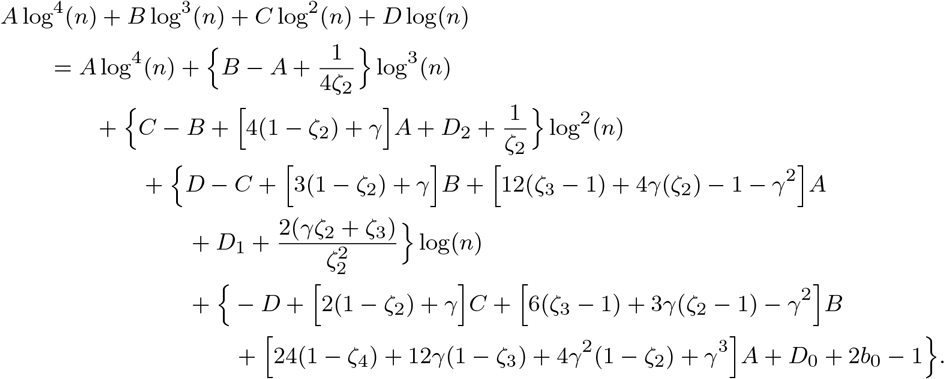

Solving the linear system implied above, we find that

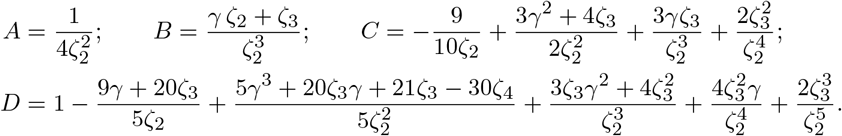

The *E*(*n*) defined by the above coefficients satisfies the recursion in (22) up to a remainder of O (log^−1^ *n*). However, if we define Δ(*n*) := |𝔼 (*Z*_*n*_) *− E*(*n*)|, then

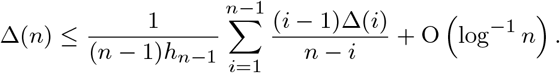

The above inequality is equivalent to the one in Aldous and Pittel (2023, equation (2.29)) in the proof of Aldous and Pittel (2023, Theorem 2.4). Repeating the arguments there, it finally follows that Δ(*n*) = O (1).

We are now ready to prove the asymptotic formulas (9)-(11) in Theorem 2.1. The proof of (9) is a straightforward consequence of (14) from Aldous and Pittel (2023, Theorem 1.2) because, due to the exchangeability of (*L*_*n,i*_):

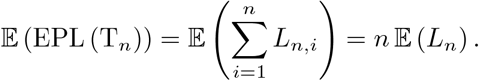

On the other hand, again due to exchangeability,

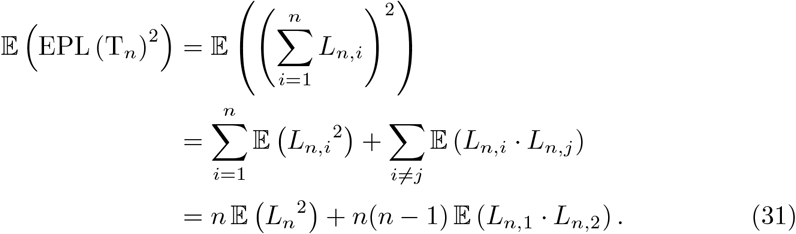

We may estimate the first term in the last identity above using (14)–(15) as follows:

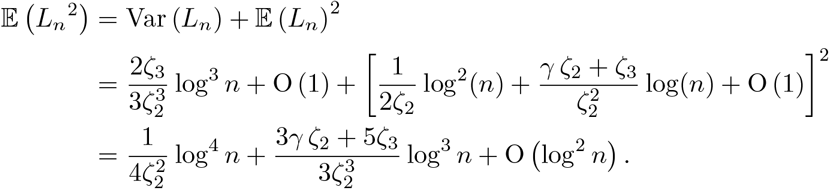

By substituting the latter expression and the asymptotic formula in (26) into (31), equation (10) follows. Finally, the formula in (11) follows from

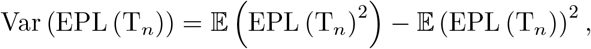

after substituting in (9)-(10), which completes the proof of Theorem 2.1.

## Appendix A Technical Lemmas for Theorem 2.1

### A.1 Proof of Lemma 5.1

To prove the lemma, we require the following three auxiliary results.

#### Lemma A.1.

*For all x* ≥ 0:

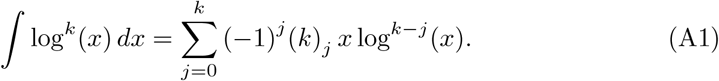

*Proof* Letting *I*_*k*_ (*x*) =∫ log^*k*^ (*x*) *dx*, the following inhomogeneous linear recursion results from integration by parts:

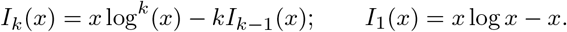

The result then follows by solving the recursion.

#### Lemma A.2.

*For all x* ≥ 0:

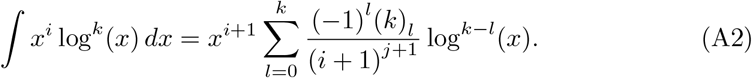

*Proof* Define *I*_*k*_ = ∫ *x*^*i*^ log^*k*^ (*x*) *dx*. Integration by parts yields the following recursion for *I*_*k*_ :

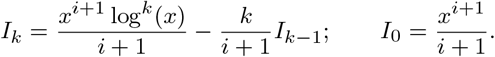

It is straightforward to then show that (A2) is the solution.

#### Lemma A.3.

*For all* 0 ≤ *x* ≤ 1:

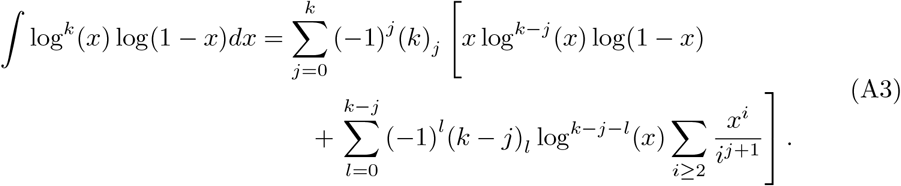

*Proof* Define *I*_*k*_ (*x*) = ∫ log^*k*^ (*x*) *dx*. Applying Lemma A.1 and integration by parts:

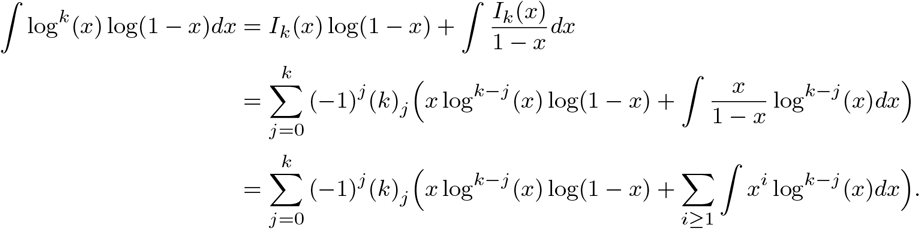

The rightmost term we can be rewritten using Lemma A.2 as follows:

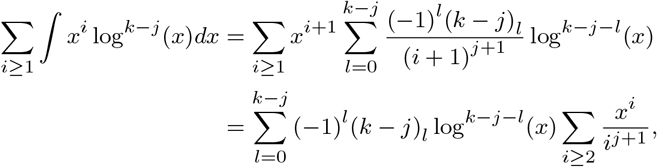

from which the lemma follows.

To prove Lemma 5.1, we start from Lemma A.3:

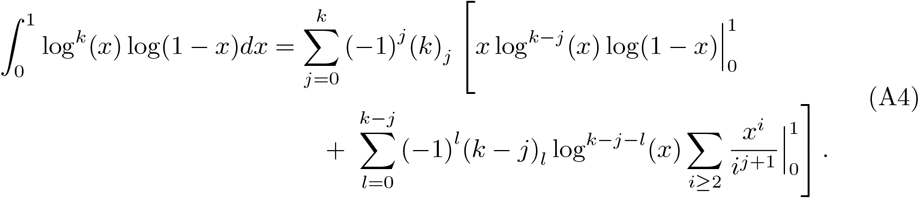

But note that for 0 ≤ *j* ≤ *k*

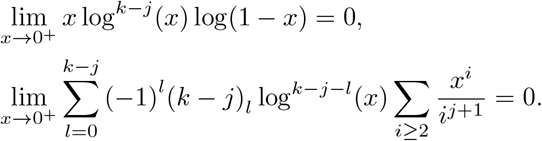

Moreover, if *j* ≠ *k* then

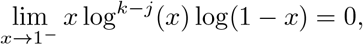

and, separately, if *l < k* − *j* then

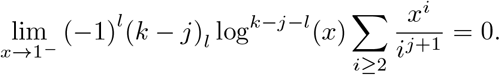

Therefore, in (A4), we need only carefully address the limit as *x* → 1^−^ in the first term when *j* = *k*, and the summation that follows when *l* = (*k* − *j*). For the latter when *j* ≠ 0, we find that

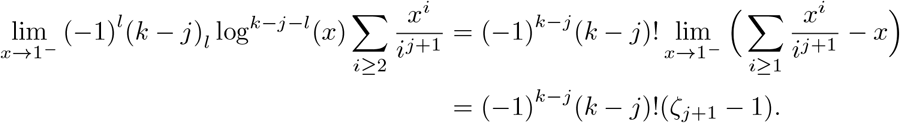

Accordingly, we can rewrite the identity in (A4) as follows:

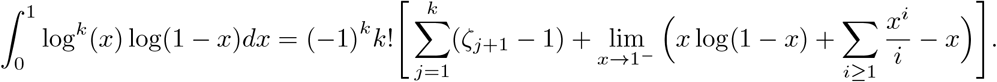

Finally, using the well-known Taylor expansion of log(1 − *x*) about *x* = 0, we find that

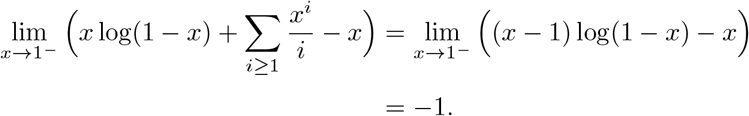

The result follows.

### A.2 Proof of Lemma 5.2

Using (17), we find

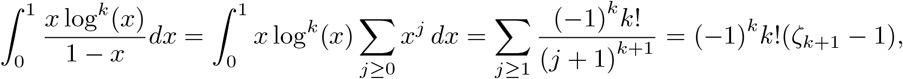

which is the desired result.

### A.3 Proof of Lemma 5.3

Using the asymptotic expansion for harmonic numbers (Graham et al. 1994, Table 452):

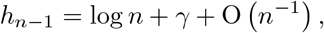

we find that (*n* − *c*)*h*_*n*−1_ = *n* log *n* + *nγ* − log *n* + O (1). Hence, for any *m >* 0:

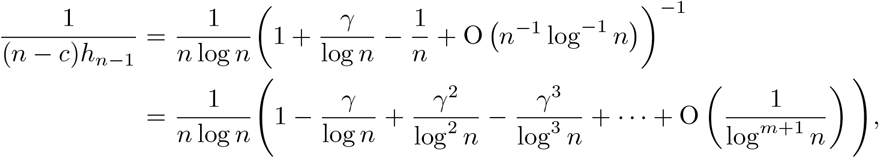

which shows the lemma.

